# Antibody-dependent priming of spontaneous germinal centers by autoreactive B cells

**DOI:** 10.64898/2026.08.27.745373

**Authors:** Danny Kim, Kristy Chiang, Andrea D. Largent, Ragan A. Pitner, Sivasankaran Munusamy Ponnan, Brock J. McKinney, Richard G. James, David J. Rawlings, Shaun W. Jackson

## Abstract

Autoreactive germinal centers (GCs) are central to autoimmune pathogenesis, yet the mechanisms by which single autoreactive B cell clones prime systemic autoimmunity remain unclear. Using the 564Igi mixed chimera model, we demonstrate that autoreactive 564Igi B cells break tolerance in wild-type B cells through an unexpected mechanism independent of cognate T cell interactions. While B cell-intrinsic TLR7 signaling was essential for spontaneous GC formation, deletion of MHC class II, CD40, or CD80/86 on GC-priming 564Igi B cells failed to prevent GCs. Instead, CRISPR-mediated deletion of *Prdm1* (encoding BLIMP-1) in 564Igi B cells ablated spontaneous GCs, implicating autoantibody production as the primary driver. These findings reveal that autoantibodies can initiate feed-forward mechanisms that propagate systemic autoimmunity, independent of B cell-intrinsic antigen presentation.

## Introduction

Systemic lupus erythematosus (SLE) and other humoral autoimmune diseases are characterized by the production of class-switched autoantibodies targeting diverse self-antigens. These autoantibodies can arise from two distinct pathways: germinal centers (GCs), specialized anatomic structures where high-affinity plasma cells and memory B cells develop through somatic hypermutation and affinity maturation; and extraofollicular (EF) B cell activation. Despite the evolution of overlapping tolerance mechanisms that normally prevent autoreactive B cell activation within GCs (1–5), spontaneous germinal centers (Spt-GCs) are aberrantly expanded in both autoimmune patients and multiple animal models of SLE (6, 7). These data implicate Spt-GCs as a critical source of pathogenic autoantibodies in autoimmune disease.

Independent animal studies have been critical for dissecting the B cell-intrinsic signals required for Spt-GC formation and the breakdown of peripheral tolerance (8, 9). These studies have shown that autoreactive B cells can initiate breaks in CD4^+^ T cell tolerance, driving T follicular helper (Tfh) cell differentiation and subsequent Spt-GC formation. B cell-intrinsic Toll-like receptor (TLR) signaling, particularly through TLR7, is critical for initiating autoreactive B cell activation (10–12). Subsequently, autoreactive B cells promote GC formation through cognate antigen presentation to CD4^+^ T cells (13, 14), provision of costimulatory signals (15), and production of inflammatory cytokines (16). However, genetic knockout models are poorly suited to distinguish between the signals required for GC initiation versus maintenance, as these approaches ablate gene function throughout the disease course. As such, understanding how single autoreactive clones initiate systemic autoimmunity remains a fundamental challenge.

The 564-immunoglobulin transgenic (564Igi) mouse provides a unique model to address this question. This strain expresses a knockin B cell receptor (BCR) specific for nucleic acid-containing antigens derived from autoimmune-prone SWRxNZB (SNF1) mice (17). To study the mechanisms underlying humoral autoimmunity, Degn et al. reconstituted irradiated wild-type (WT) recipients with different ratios of 564Igi and WT bone marrow (BM). Resulting chimeras subsequently developed Spt-GCs that were proportional in size to the input 564Igi percentage. Remarkably, whereas autoreactive 564Igi B cells were required to initiate GC formation, the majority of B cells within established GCs were WT-derived, and WT B cells were sufficient to propagate self-perpetuating GCs (18). This paradox suggests that 564Igi B cells prime breaks in WT B cell tolerance through as yet undefined mechanisms. As such, the 564Igi mixed chimera model provides a novel strategy to genetically interrogate the B cell-intrinsic signals required for the autoimmune GC initiation vs. maintenance.

Here, we systematically dissected the cell-intrinsic requirements for induction of breaches in tolerance by an individual autoreactive B cell clone. Our findings challenge the prevailing model that B cells function primarily as antigen-presenting cells in autoimmunity initiation, instead revealing autoantibodies as critical initial mediators of systemic tolerance breakdown.

## Results

### Initiation of GC formation by 564Igi B cells requires B cell-intrinsic TLR7 signaling

Toll-like receptor 7 (TLR7) is a pattern-recognition receptor activated by RNA that synergizes with B cell receptor signals to break tolerance to RNA-associated autoantigens during lupus pathogenesis (19). 564Igi-derived antibodies exhibit polyclonal reactivity against single-stranded DNA (ssDNA), ssRNA, and related RNA-associated autoantigens, and TLR7 deletion prevents autoantibody production in homozygous 564Igi mice (17). Previous studies using mixed BM chimeras revealed that TLR7 expression by WT B cells is necessary to sustain ongoing GC reactions initiated by 564Igi B cells. However, this work did not address the B cell-intrinsic requirement for TLR7 in the initial breach of tolerance induced by 564Igi B cells.

To isolate this initiating step, we generated mixed BM chimeras comprising 2:1 ratios of congenically-marked WT (CD45.1) and *Tlr7*-sufficient or -deficient 564Igi B cells (CD45.2). Briefly, CD45.1^+^ recipient mice were lethally irradiated (450 cGy x 2) and reconstituted with 6 x 10^6^ bone marrow (BM) cells comprising the following groups: i) CD45.2^+^ WT (1 part) plus CD45.1^+^ WT (2 parts); ii) *Tlr7^+/+^.*564Igi (1 part; CD45.2^+^) plus WT (2 parts: CD45.1^+^); iii) *Tlr7^-/-^.*564Igi (1 part; CD45.2^+^) plus WT (2 parts: CD45.1^+^) (**Fig. 1A**). Resulting chimeras were sacrificed at 6 weeks post BM transplant for quantification of Spt-GCs. In each of the 564Igi chimera models, homozygous 564 heavy and light chain expression was confirmed via anti-idiotype staining of CD45.2⁺ 564Igi B cells in respective chimera models. Furthermore, CD45.2⁺ 564Igi B cells displayed expected developmental arrest at the transitional stage, a hallmark of autoreactive B cells undergoing negative selection in polyclonal settings (**Fig. 1B**).

**Figure 1:**
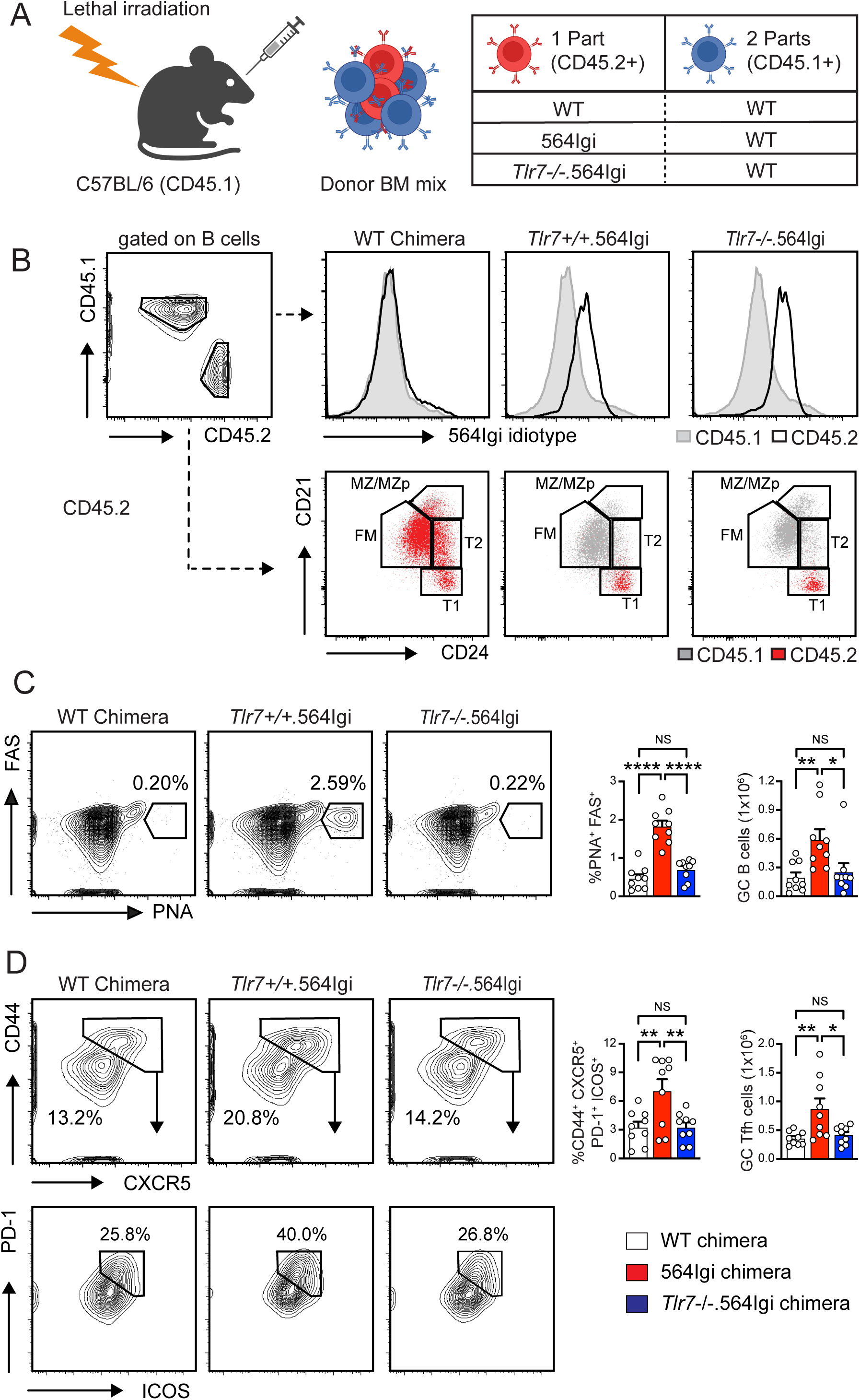
TLR7-dependent initiation of spontaneous GCs by 564Igi B cells. **(A)** Schematic overview of experimental design of *Tlr7*-deficient 564Igi BM chimera models. Lethally irradiated wild-type C57BL/6 (WT; CD45.1⁺) recipient mice were reconstituted with a 2:1 mixture of wild-type (WT; CD45.1⁺) BM and donor (CD45.2⁺) BM from one of the following strains: WT, *Tlr7^+/+^.*564Igi, and *Tlr7^-/-^.*564Igi. **(B)** Representative flow cytometry plots showing the gating strategy 6 weeks post-reconstitution. Upper right panels: donor-derived (CD45.2⁺) B cells were analyzed for expression of 564Igi by anti-idiotype staining. Gray shaded histogram indicates CD45.1^+^ WT B cells. Lower right panels: In both *Tlr7^+/+^.*564Igi, and *Tlr7^-/-^.*564Igi chimeras, CD45.2⁺ 564Igi B cells (red) exhibit developmental arrest at the CD21^low^CD24^high^ transitional 1 (T1) stage. CD45.1^+^ WT B cells shaded gray. B cell subset abbreviations: T1, transitional 1; T2, transitional 2, FM, follicular mature; MZ/MZp, marginal zone/marginal zone precursor. **(C)** Representative flow cytometry plots (left) and quantification (right) of splenic PNA⁺FAS⁺ germinal center (GC) B cells. Left panel: gated on B220^+^CD19^+^ B cells. Number indicates percentage within gate. **(D)** Representative flow cytometry plots (left) and quantification (right) of splenic CD44⁺CXCR5⁺PD-1⁺ICOS⁺ GC T follicular helper (Tfh) cells. Left: upper panels gated on CD4^+^ T cells; lower panels gated on CD44⁺CXCR5⁺ T cells. Number indicates percentage within each gate. **(C, D)** Data representative of ≥3 independent BM chimera experiments. Bar graphs indicate mean ± SEM. Each symbol indicates an individual WT (white), *Tlr7^+/+^.*564Igi (red), and *Tlr7^-/-^.*564Igi (blue) animal. *, *P*<0.05; **, *P*<0.01; ****, *P*<0.0001, by one-way ANOVA with Tukey’s multiple comparison test. NS, not significant.

Consistent with previous studies (18), chimeras reconstituted with *Tlr7^+/+^.*564Igi cells developed robust splenic GCs (**Fig. 1C**). In contrast, GC formation was abrogated in chimeras that received *Tlr7^-/-^.*564Igi cells. This loss of GCs corresponded with a profound reduction in splenic CD4^+^ germinal center T follicular helper cells (GC Tfh; CD44^+^CXCR5^+^PD1^+^ICOS^+^) (**Fig. 1D**). Together, these data demonstrate that B cell-intrinsic TLR7 signaling by autoreactive 564Igi B cells is the critical initiating event that breaks tolerance and primes subsequent Spt-GC and Tfh cell expansion.

### Autoreactive B cells can prime spontaneous GCs without classical cognate T cell interactions

The ability of B cells to capture and present specific antigens via their B cell receptor is a critical driver of humoral autoimmunity (13, 14, 20). This function is thought to propagate disease by focusing T cell help on self-antigens, leading to a breach of tolerance. While established autoreactive GCs depend on this B cell:T cell crosstalk, it remains undefined whether B cell antigen presentation or costimulation is required to initiate tolerance breaks. We therefore sought to determine if these canonical pathways are necessary for 564Igi B cells to prime the Spt-GC response.

To test roles for antigen presentation and costimulation, we generated parallel sets of mixed bone marrow (BM) chimeras. Lethally irradiated CD45.1⁺ recipient mice were reconstituted with a 2:1 mixture of WT (CD45.1⁺) BM and donor (CD45.2⁺) BM. The experimental donor populations consisted of 564Igi B cells that were additionally deficient in MHCII (*MhcII^-/-^*), CD40 (*Cd40^-/-^*), or both CD80 and CD86 (*Cd80^-/-^*.*Cd86^-/-^*). Control chimeras received either WT (CD45.2⁺) or unmodified 564Igi donor BM (**Fig. 2A**). As in the *Tlr7* chimera model, CD45.2^+^ 564Igi B cells across all genotypes displayed developmental arrest at the transitional 1 (T1) stage (**Fig. 2B**). We also confirmed the absence of MHCII and CD40 expression on the respective CD45.2⁺ 564Igi donor B cell populations (**Fig. 2C**).

**Figure 2:**
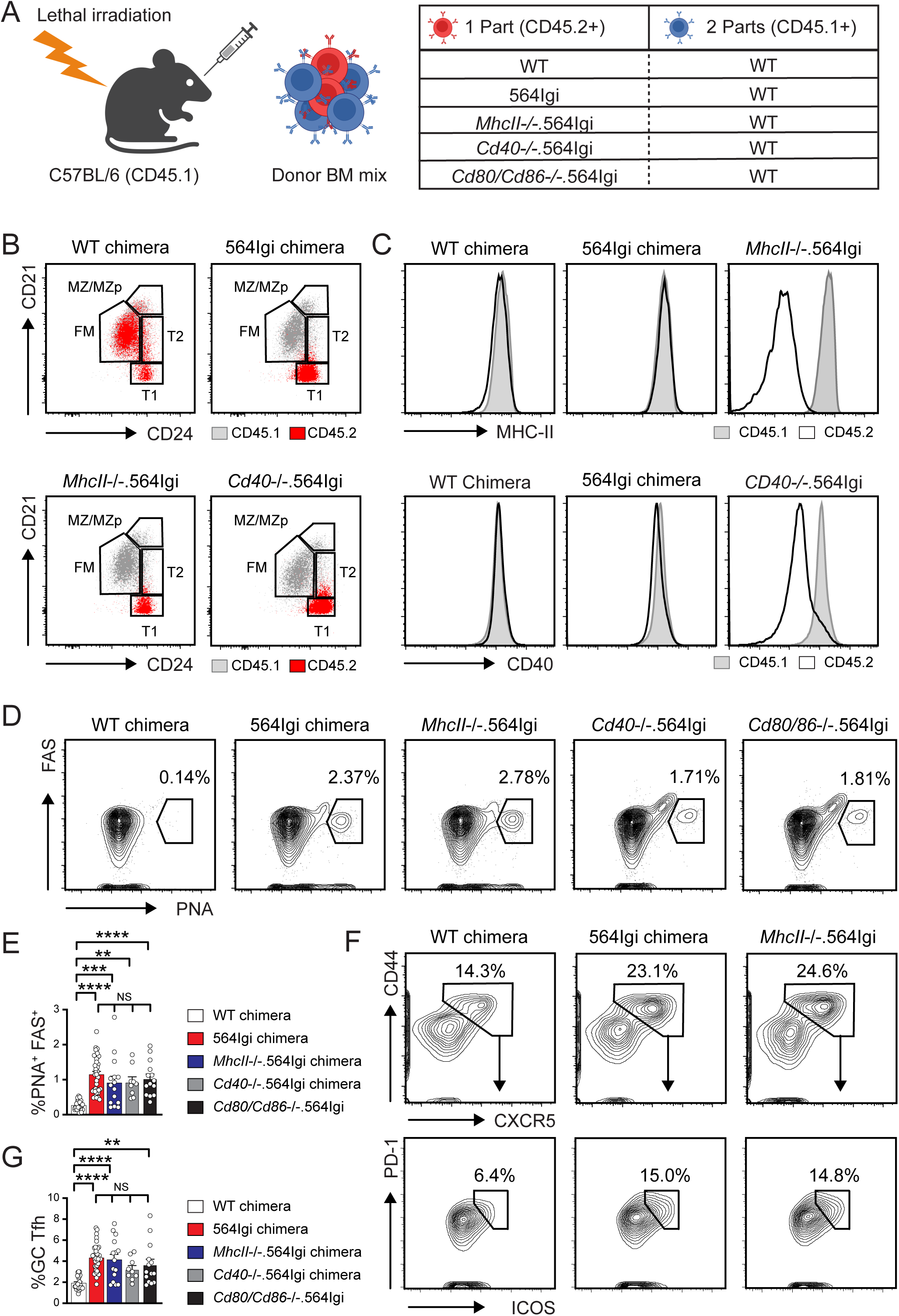
564Igi B cells can initiate spontaneous GCs independent of cognate CD4^+^ T cell interactions. **(A)** Schematic overview of experimental design. Lethally irradiated CD45.1⁺ WT mice were reconstituted with 2 parts CD45.1⁺ WT BM and 1 part CD45.2⁺ WT, 564Igi, *MhcII^-/-^.*564Igi, *Cd40^-/-^.*564Igi, or *Cd80^-/-^.Cd86^-/-^.*564Igi donor BM. **(B)** Representative flow plots showing developmental arrest of CD45.2⁺ 564Igi B cells (red) at the CD21^low^CD24^high^ transitional 1 (T1) stage in indicated chimeras. CD45.1^+^ WT B cells shaded gray. B cell subset abbreviations: T1, transitional 1; T2, transitional 2, FM, follicular mature; MZ/MZp, marginal zone/marginal zone precursor. **(C)** Histograms showing MHCII (upper panels) and CD40 (lower panels) expression on total CD45.1^+^ vs. CD45.2^+^ B cells in indicated chimera models. CD45.1^+^, gray shaded histogram vs. CD45.2^+^, solid black line. CD80/CD86 expression on 564Igi B cells not shown since these markers were omitted from the flow cytometry panels. **(D)** Representative flow cytometry plots showing splenic PNA⁺FAS⁺ GC B cells in respective chimera models. Gated on B220^+^CD19^+^ B cells. Number indicates percentage within gate. **(E)** Quantification of splenic PNA⁺FAS⁺ GC B cells (% of total CD19^+^B220^+^ B cells). **(F)** Representative flow cytometry plots (gated on splenic CD4^+^ T cells) showing identification of CD44⁺CXCR5⁺PD-1⁺ICOS⁺ GC T follicular helper (Tfh) cells. Number indicates percentage within each gate. **(G)** Quantification of splenic CD44⁺CXCR5⁺PD-1⁺ICOS⁺ GC T follicular helper (Tfh) cells (% of total CD4^+^ T cells). **(E, G)** Bar graphs indicate mean ± SEM. Each symbol indicates an individual WT (white), 564Igi (red), *MhcII^-/-^.*564Igi (blue), *Cd40^-/-^.*564Igi (gray), or *Cd80^-/-^.Cd86^-/-^.*564Igi (black) animal. *, *P*<0.05; **, *P*<0.01; ***, *P*<0.001; ****, *P*<0.0001, by one-way ANOVA with Tukey’s multiple comparison test. NS, not significant. Data representative of five WT, seven 564Igi, three *MhcII^-/-^.*564Igi, two *Cd40^-/-^.*564Igi, and two *Cd80^-/-^.Cd86^-/-^.*564Igi independent competitive chimeras.

Based on the central role of these canonical antigen presentation pathways in adaptive immunity, we hypothesized that deleting any of these molecules from the 564Igi B cells would abrogate their ability to prime GC and Tfh cell formation. Contrary to our hypothesis, B cell-intrinsic antigen presentation and costimulation were completely dispensable for the initiation of the autoimmune response. Chimeras receiving 564Igi B cells lacking MHCII, CD40, or CD80/86 all developed robust splenic GCs, with frequencies comparable to the positive control 564Igi chimeras (**Fig. 2D, E**). Likewise, the expansion of the CD44^+^CXCR5^+^PD1^+^ICOS^+^ GC Tfh cells was not hindered by the absence of these molecules on the initiating autoreactive B cells (**Fig. 2F, G**). In all experimental groups, both the GC and Tfh responses were significantly greater than in the WT control chimeras. These data reveal a striking divergence between the requirements for initiating versus sustaining an autoimmune response, showing that the initial break in tolerance by highly autoreactive 564Igi B cells is independent of antigen presentation and costimulatory signals.

### B cell-intrinsic IRF5 and IL-6 are dispensable for initiating spontaneous GCs

The finding that 564Igi B cells can prime autoimmunity independently of their canonical antigen presentation and costimulatory functions led us to investigate the critical B cell-intrinsic pathways downstream of TLR7. We focused on the transcription factor IRF5 and its downstream effector cytokine, interleukin-6 (IL-6), which form a well-established pro-inflammatory axis in humoral autoimmunity. Indeed, our prior work demonstrated that B cell-derived IL-6 promotes Tfh cell differentiation and Spt-GC formation (16). Given that IRF5 promotes B cell IL-6 production and orchestrates other pro-autoimmune B cell functions (21–23), we hypothesized that this IRF5/IL-6 pathway would be the essential link between TLR7 sensing and the initiation of the Spt-GC response.

To test this hypothesis, we generated mixed bone marrow chimeras by reconstituting lethally irradiated CD45.1⁺ recipients with a 2:1 mixture of WT BM and donor BM from either *Il6^-/-^*.564Igi or *Irf5^-/-^*.564Igi mice, alongside appropriate WT and 564Igi controls (**Fig. 3A**). We anticipated that, similar to TLR7 deficiency, the absence of either IRF5 or IL-6 in the initiating 564Igi B cells would abrogate Spt-GC formation.

**Figure 3:**
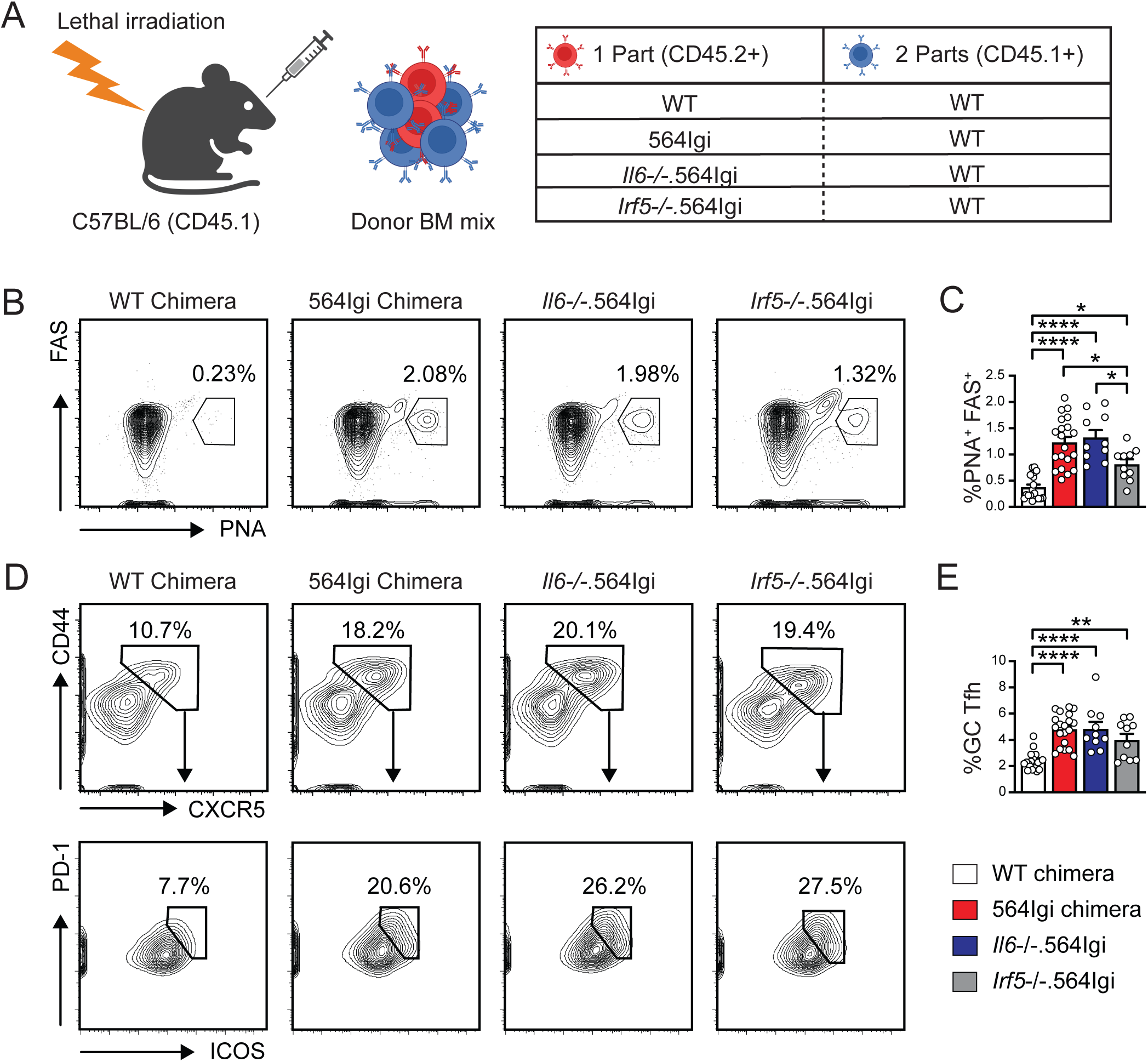
IL-6 produced by autoreactive 564Igi B cells is not required to initiate spontaneous GCs. **(A)** Schematic overview of experimental design. Lethally irradiated CD45.1⁺ WT mice were reconstituted with 2 parts CD45.1⁺ WT BM and 1 part CD45.2⁺ WT, 564Igi, *Il6^-/-^.*564Igi, or *Irf5^-/-^.*564Igi donor BM. **(B)** Representative flow cytometry plots showing splenic PNA⁺FAS⁺ GC B cells in respective chimera models. Gated on B220^+^CD19^+^ B cells. Number indicates percentage within gate. **(C)** Quantification of splenic PNA⁺FAS⁺ GC B cells (% of total CD19^+^B220^+^ B cells). **(D)** Representative flow cytometry plots (gated on splenic CD4^+^ T cells) showing identification of CD44⁺CXCR5⁺PD-1⁺ICOS⁺ GC T follicular helper (Tfh) cells. Number indicates percentage within each gate. **(E)** Quantification of splenic CD44⁺CXCR5⁺PD-1⁺ICOS⁺ GC T follicular helper (Tfh) cells (% of total CD4^+^ T cells). **(C, E)** Bar graphs indicate mean ± SEM. Each symbol indicates an individual WT (white), 564Igi (red), *Il6^-/-^.*564Igi (blue), or *Irf5^-/-^.*564Igi (gray) animal. *, *P*<0.05; **, *P*<0.01; ****, *P*<0.0001, by one-way ANOVA with Tukey’s multiple comparison test. Data representative of four WT, four 564Igi, two *Il6^-/-^.*564Igi, and two *Irf5^-/-^.*564Igi independent competitive chimeras.

Strikingly, and contrary to our hypothesis, the data revealed that both the IRF5 and IL-6 pathways were dispensable for the initiation of autoimmunity. Six weeks after reconstitution, analysis of the spleens showed that chimeras receiving either *Il6^-/-^*.564Igi or *Irf5^-/-^*.564Igi B cells developed robust Spt-GC responses (**Fig. 3B, C**). This pattern was mirrored in the cognate T cell response, where Tfh cell expansion was similarly unimpeded by the loss of B cell-intrinsic IL-6 or IRF5 (**Fig. 3D, E**). Although chimeras receiving *Irf5^-/-^*.564Igi cells displayed a modest reduction in GC B cells relative to 564Igi positive controls, the frequency of PNA⁺FAS⁺ GC B cells remained significantly greater than WT controls. Thus, while these data suggest a partial contribution for IRF5-dependent mechanisms in Spt-GC initiation, the primary conclusion remains that, like the redundant roles for B cell antigen presentation, neither IL-6 production by the autoreactive B cell nor its broader IRF5-dependent functions are essential for priming the autoimmune cascade.

### Autoantibody secretion by priming autoreactive B cells is required to initiate spontaneous GC formation

The above findings revealed that canonical B cell effector functions, including antigen presentation, are surprisingly dispensable for initiating autoimmunity in the 564Igi model. This prompted us to investigate an alternative hypothesis that the critical initiating signal is delivered not through direct cell-to-cell contact, but through the secretion of autoantibodies. We posited that autoantibodies locally produced by 564Igi B cells form pathogenic immune complexes that are then captured and presented by other antigen-presenting cells, including follicular dendritic cells (FDC), to prime the T cell response. Although a previous study found that passive transfer of 564Igi serum was insufficient to induce Spt-GCs (20), we reasoned that this approach may have failed to achieve the necessary local concentration or sustained titer of autoantibodies required to break tolerance.

To definitively test the requirement for antibody secretion, we sought to uncouple the presence of the 564Igi B cell from its ability to differentiate into an antibody-secreting plasma cell via deletion of *Prdm1*, the gene encoding the master transcriptional regulator of plasma cell differentiation, BLIMP-1 (24). Since *Prdm1^-/-^* mice were unavailable to intercross with 564Igi mice, we employed a CRISPR-Cas9 gene-editing approach to knock out *Prdm1* in mature 564Igi B cells prior to adoptive transfer. We previously showed that this approach achieves >90% biallelic *Prdm1* knockout rates in mature B cells, resulting in near complete loss of plasma cell differentiation and antibody production in vitro (25).

Because this CRISPR-based strategy uses mature B cells, not BM stem cells, we first had to validate that the adoptive transfer of activated B cells can effectively prime Spt-GCs. To do this, lethally irradiated CD45.1^+^ recipient mice were reconstituted with CD45.1^+^ WT BM plus varying doses of in vitro-activated 564Igi B cells. As early as 1 week after cell transfer, 564Igi-derived anti-Sm/RNP IgG2a autoantibodies could be detected in recipient animals with titers sustained for 6 weeks (**Fig. 4A**). In addition, this experiment confirmed that the adoptive transfer of activated 564Igi B cells was sufficient to initiate Spt-GC formation, with the resulting GCs being predominantly composed of CD45.1^+^ WT-derived B cells, thus recapitulating the key features of the original model (**Fig. 4B-D**). Having established that a transfer of 5 x 10^6^ cells was sufficient for GC priming, we selected this dose for subsequent CRISPR experiments.

**Figure 4:**
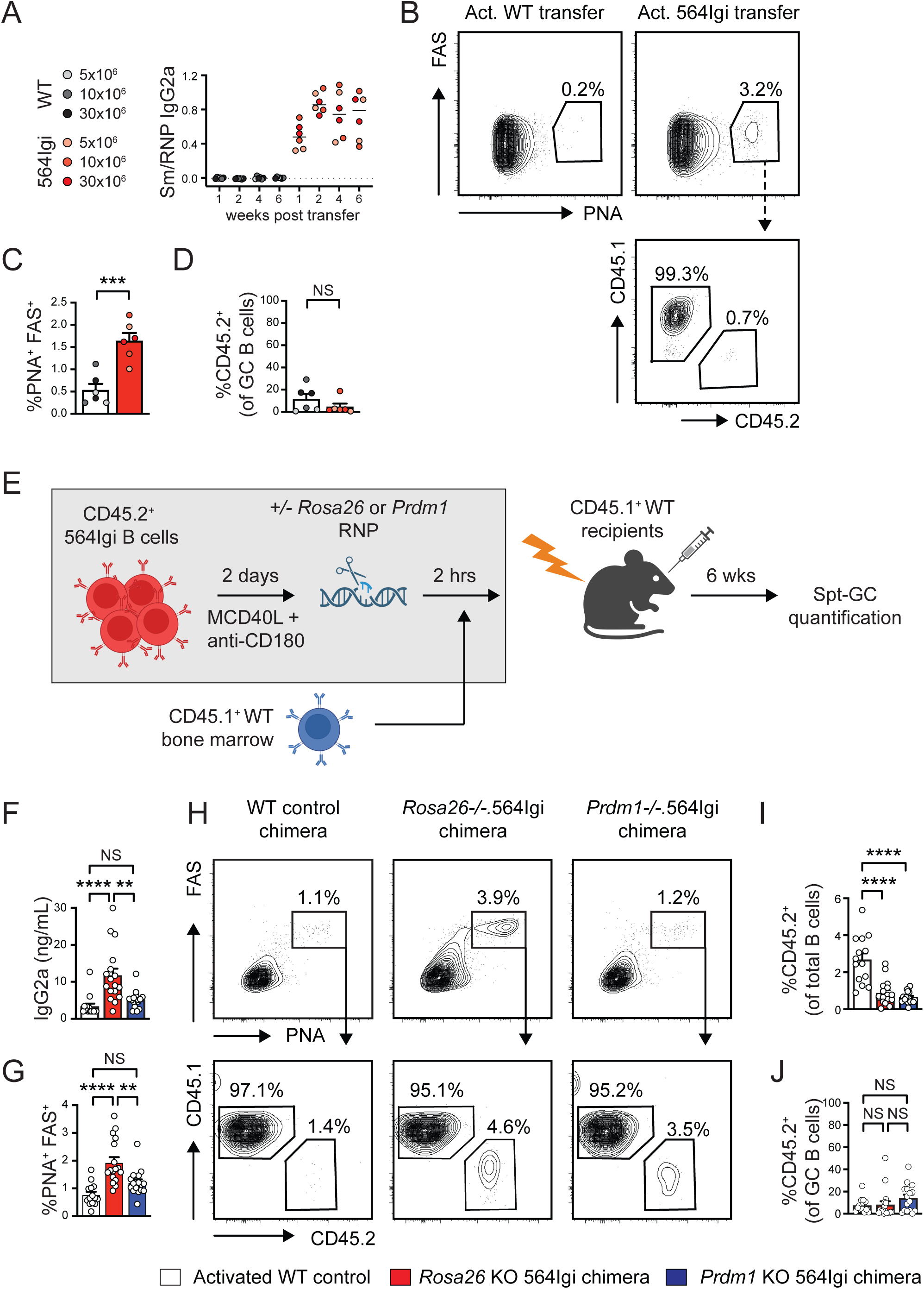
Antibody-dependent priming of spontaneous GCs by 564Igi B cells. **(A-D)** <u>Activated 564Igi B cell transfer experiment:</u> **(A)** Anti-SmRNP IgG2a titers in recipients of WT or 564Igi activated B cells at indicated weeks after B cell transfer. **(B)** Representative flow cytometry plots showing expansion of PNA⁺FAS⁺ GC B cells in chimera receiving 10 x 10^6^ activated 564Igi B cells (upper panels). Lower panel shows CD45.1⁺ vs. CD45.2⁺ proportions within the GC gate of a representative 564Igi B cell recipient. Upper panels: Gated on B220^+^CD19^+^ B cells. Lower panel: Gated on PNA⁺FAS⁺ GC B cells. Number indicates percentage within gate. **(C)** Quantification of splenic PNA⁺FAS⁺ GC B cells (% of total CD19^+^B220^+^ B cells). **(D)** % CD45.2⁺ as a proportion of splenic PNA⁺FAS⁺ GC B cells. **(A, C, D)** Graphs indicate mean ± SEM. Each symbol indicates an individual recipient of activated WT (grey) or 564Igi (red) B cells, color coded by transfer B cell number. ***, *P*<0.001; NS, not significant, by unpaired Student *t* test. **(E-J)** <u>CRISPR-edited B cell transfer experiment:</u> **(E)** Schematic overview of experimental design. CD45.2⁺ 564Igi B cells were activated for 2 days using murine multimeric CD40L and anti-CD180, then electroporated with Cas9 ribonucleoproteins (RNPs) targeting either *Prdm1* or control locus *Rosa26*. CRISPR-edited B cells (1 part) and CD45.1⁺ WT BM (2 parts) were adoptively transferred into lethally irradiated CD45.1⁺ WT mice and splenic GCs quantified at 6 weeks post transfer. **(F)** Total serum IgG2a titers in respective chimera models. **(G)** Quantification of splenic PNA⁺FAS⁺ GC B cells (% of total CD19^+^B220^+^ B cells). **(H)** Representative flow cytometry plots showing PNA⁺FAS⁺ GC B cells in indicated experimental models (upper panels; gated on B220^+^CD19^+^ B cells), and CD45.1⁺ vs. CD45.2⁺ proportions within GCs (lower panels). Number indicates percentage within gate. **(I, J)** % CD45.2⁺ B cells as a proportion of total B cells **(I)** and GC B cells **(J)** in indicated experimental models. **(F, G, I, J)** Bar graphs indicate mean ± SEM. Each symbol indicates an individual recipient of activated WT B cells (white), *Rosa26*-edited 564Igi B cells (red), or *Prdm1*-edited 564Igi B cells (blue). **, *P*<0.01; ****, *P*<0.0001; NS, not significant, by one-way ANOVA with Tukey’s multiple comparison test. Data representative of four independent CRISPR-edited B cell transfer experiments.

For the definitive experiment, we employed CRISPR-Cas9 gene editing to delete *Prdm1* in 564Igi B cells prior to adoptive transfer. Briefly, splenic B cells were activated with murine multimeric CD40L and anti-CD180, cultured for 2 days, then electroporated with ribonucleoprotein complexes (RNPs) consisting of Cas9 bound to single guide RNA (sgRNA) targeting either *Prdm1* or a control locus (*Rosa26*). After 2 hours rest, 5×10^6^ *Prdm1-* or *Rosa26-*edited B cells were adoptively transferred into lethally-irradiated CD45.1^+^ recipients that had been reconstituted with 6 x 10^6^ CD45.1^+^ WT bone marrow (BM). In vitro-activated CD45.2^+^ WT B cells without gene editing were used to generate WT control chimeras (**Fig. 4E**).

As hypothesized, *Prdm1* deletion in transferred 564Igi B cells resulted in reduced 564Igi-derived IgG2a serum antibodies in the recipient mice that was accompanied by a significant decrease in the formation of Spt-GCs (**Fig. 4F-H**). Importantly, the lack of GC formation could not be attributed to poor survival of the edited cells, as we observed no difference in the persistence of CD45.2^+^ *Prdm1*-edited 564Igi B cells compared to the *Rosa26*-edited controls (**Fig. 4I**). This finding is consistent with the known role of BLIMP-1 in directing plasma cell differentiation, not B cell survival (24). In addition, we previously showed that Blimp1 deletion can result in antigen-specific *Prdm1*-knockout B cells outcompeting endogenous B cells for GC entry (25). However, we observed no such expansion of CD45.2^+^ Blimp1-knockout 564Igi B cells, relative to *Rosa26*-edited controls, within GCs of *Prdm1-*edited BM chimeras (**Fig. 4H, J**). In summary, these data demonstrate that blocking the ability of 564Igi B cells to differentiate into antibody-secreting cells largely prevents the initiation of Spt-GCs.

## Discussion

Spontaneous, autoreactive GCs are central to the pathogenesis of SLE and contribute to the diversification of the autoantibody repertoire through epitope spreading (6, 7, 18). Understanding the mechanisms that break B cell tolerance and trigger the formation of these Spt-GCs therefore has significant clinical implications. While studies in murine lupus models have uncovered a central role for dysregulated B cell signaling and B-T cell crosstalk in this process (26–28), a key technical challenge has remained. Since humoral autoimmunity develops stochastically without a clear starting point, it has been challenging to distinguish the B cell-intrinsic signals that initiate activation from those that sustain Spt-GC responses. In this context the 564Igi BM chimera model, in which a single autoreactive B cell clone primes a self-sustaining, polyclonal GC response, provides a unique opportunity to genetically interrogate the distinct signals required for the initiation versus the maintenance of humoral autoimmunity (18).

While our data confirm the expected role of TLR7 signaling in initiating autoreactive B cell activation, we surprisingly found that GC priming can occur independently of the initiating B cells’ capacity to present self-antigens or engage cognate T cell help. Instead, we reveal that feed-forward autoantibody production plays a critical, non-redundant role in priming this breach of tolerance. We propose that crosslinking of the 564Igi B cell receptor by RNA-containing self-antigens initiates TLR7-dependent and T cell-independent plasma cell differentiation. Although not tested here, we hypothesize that resulting autoantibodies could be captured by follicular dendritic cells (FDC) as immune complexes (IC) to prime GCs. Our study is thus consistent with prior work showing that FDC capture of circulating IC promotes TLR7-dependent activation and IFN-α production (29), and offers a plausible mechanism for the genesis of the IC that initiate this FDC-driven amplification of the autoimmune response.

An important caveat to this proposed model is that *Prdm1* deletion exerts impacts on B cells beyond blocking plasma cell differentiation. BLIMP-1 functions as a transcriptional repressor, silencing the expression of core B cell genes including *Pax5* and *Bcl6*, to drive the transition from GC B cell to antibody-secreting plasma cell. As such, *Prdm1* deletion not only limits antibody secretion but also impairs GC exit, expands the memory B cell pool, and promotes B cell antigen presentation by maintaining MHC Class II and costimulatory molecule expression (30–33). In principle, these secondary effects could influence Spt-GC formation independently of autoantibody production.

However, we believe that these plasma cell-independent effects of *Prdm1* deletion are unlikely to account for the observed loss of Spt-GCs for several reasons. First, *Prdm1* deletion is limited to priming CD45.2^+^ 564Igi B cells, whereas the CD45.1^+^ WT B cells that comprise the majority of Spt-GCs retain BLIMP-1. Second, although *Prdm1* deficiency would be expected to enhance antigen presentation to cognate CD4^+^ T cells, we show that these B cell functions are redundant for Spt-GC priming by autoreactive 564Igi B cells (**Fig. 2**). Taken together, our data support a model in which autoantibody secretion, rather than ancillary *Prdm1* functions, are critical for Spt-GC priming in the 564Igi chimera model.

Importantly, our findings do not conflict with recent data showing that B cells drive MHC-dependent epitope spreading within autoimmune GCs (20). A key distinction is that previous work focuses on the propagation of the autoimmune response within the GC, whereas our current study addresses the initial priming events that leads to GC formation itself. We find that this initial step is antibody-dependent and does not require antigen presentation by the initiating B cell clone. It is therefore plausible that the antibody-driven mechanism we describe here primes autoimmune GCs, providing the microenvironment for ongoing B-T cell interactions and MHC-dependent epitope spreading detailed by Fahlquist-Hagert, et al (20).

A limitation of our study is that we focused on the mechanisms underlying Spt-GC formation, without addressing potential impacts on extrafollicular (EF) B cell activation pathways. We acknowledge that EF B cell responses make an important, parallel contribution to autoantibody production in SLE (34–36). However, the 564Igi mixed chimera model is uniquely suited to interrogating GC dynamics, since 564Igi and WT B cells exert separate contributions to Spt-GC priming and maintenance, providing a clear cellular readout of breaks in immune tolerance. Whether the B cell-intrinsic signals required to prime humoral autoimmunity differ between EF and GC pathways, and whether autoantibody-dependent priming extends to EF B cell activation, remain open questions for future study.

In summary, our study advances the understanding of the early events in autoimmune GC formation by identifying a new mechanisms by which autoreactive B cell clones can promote breaks in immune tolerance during the pathogenesis of humoral autoimmunity.

## Materials and Methods

### Sex as a biological variable

All CD45.1^+^ recipient mice and CD45.1^+^ donor BM used in this study were male. CD45.2^+^ donor BM (WT, 564Igi, and respective genetic intercrosses) was derived from both male and female mice, depending on availability of each strain. Within each experimental cohort, sex was matched between donor and recipient populations to control for potential sex-based effects on chimerism and immune reconstitution.

### Mice

Animal strains include: B6 (C57BL/6J; Jax stock #000664), B6 CD45.1 (Jax stock #002014), *Tlr7^-/-^* (Jax stock #008380; (37)), *Il6^-/-^* (Jax stock #002650; (38)), *MhcII^-/-^* (Jax stock #003584; (39)), *Cd80^-/-^.Cd86^-/-^* (Jax stock #003610; (40)), and *Cd40^-/-^*(Jax stock #002928; (41)). *Irf5^-/-^* mice corrected for *Dock2* mutation (42, 43) were generously provided by Dr. Michael Gale (University Washington, WA, USA) and 564Igi mice (17) were provided by Dr. Michael C. Carrol (Boston Children’s, Boston, MA). All mice were maintained on a C57BL/6 background and bred/maintained in a specific pathogen-free facility at Seattle Children’s Research Institute. Animal experiments were conducted according to the protocols approved by the Institutional Animal Care and Use Committee at Seattle Children’s Research Institute.

### Antibodies and reagents

Anti-murine antibodies used in this study include the following: Blimp-1-AlexaFluor647 (563643), CD19-PE (557399), CD19-PE-Cy7 (552854), CD24-PE (553262), CD45R/B220-FITC (553087), CD45.2-APC-Cy7 (560694), CD80-BV421 (566285), CD86-PE-Cy7 (560582), CD95(Fas)-BV421 (562633), CD95(Fas)-PE (554258), CD278(ICOS)-PE (552146), CD185(CXCR5)-Biotin (551960), Streptavidin-APC-Cy7 (554063), Streptavidin-PE-Cy7 (557598) from BD Biosciences. CD3ε-BV510 (100353), CD8a-BV510 (100751), CD8a-PE-Cy7 (100722CD19-BV605 (115540), CD21/CD35(CR2/CR1)-APC (123412), CD24-Pacific Blue (101820), CD44-APC-Cy7 (103028), CD45R/B220-BV510 (103247), CD45R/B220-BV650 (103241), CD45R/B220-PerCp-Cy5.5 (103236), CD197 (CCR7)-PerCp-Cy5.5 (120116), CD267(TACI)-PE (133403), F4/80-BV510 (123135), H-2K^d^-BV510 (116625), I-A/I-E(MHC II)-APC (107614), NK-1.1-BV510 (108737), were from BioLegend. CD38-APC (501123048), CD45R/B220-APC-eFluor780 (501129359), CD45.1-APC (501123055), CD45.1-PerCp-Cy5.5 (5015798), CD45.2-Alexa Fluor700 (5016896), CD45.2-APC (5014979), CD279(PD-1)-FITC (501128727), T-bet-PerCp-Cy5.5 (501130726) from Fisher Scientific. CD19-Alex Fluor700 (56-0193-80), BCL6-PE (12545382), IRF4-PerCp-Cy5.5 (46985882), I-A/I-E (MHC II)-Biotin (13-5321-82) from Life Technologies. CD4-APC (1540-11), was from SouthernBiotech. CD45.1-FITC (11-0453-82), CD45.1-eFluor450 (48-0453-82), NHS Ester (Live-dead dye)-AF350 (A10168) from Thermo Fisher Scientific. Peanut Agglutinin (PNA)-FITC was from Vector Labs. 564Igi Idiotype Antibody was generated from biotinylating Anti-564-Tg IgG generated and purified from hybridoma cell using Lightning-Link Rapid Type A Biotin Antibody Labeling Kit (370-0015) from Novus Biologicals. Sm-RNP is from Arotec Diagnostics.

### Bone marrow chimeras

For mixed BM chimera models, BM were harvested from the femora and tibiae of B6 (WT CD45.2), B6 CD45.1 (WT), 564Igi (CD45.2), *Tlr7^-/-^*.564Igi (CD45.2), *MhcII^-/-^*.564Igi (CD45.2), *Cd80^-/-^.Cd86^-/-^*.564Igi (CD45.2), *Cd40^-/-^*.564Igi (CD45.2), *Il6^-/-^*.564Igi (CD45.2), and *Irf5^-/-^*.564Igi (CD45.2) mice. Extracted cells were resuspended in RBC lysis buffer (ACK; Gibco, Thermo Fisher Scientific) and filtered through 40-µm strainers (Corning). CD138-depleted donor BM (564Igi and 564Igi knockout models) was mixed with CD45.1 (WT) BM at 1:2 ratio. Total of 6 × 10^6^ cells was injected retro-orbitally into lethally irradiated (450 cGy × 2 doses) B6 CD45.1 recipients. Injected recipients were placed on Baytril-medicated water for 2 weeks after irradiation. Chimera animals were sacrificed at 6 weeks to quantify splenic GCs. For activated B cell transfer models (see below), 6 × 10^6^ B6 CD45.1 (WT) BM cells were injected retro-orbitally into lethally irradiated (450 cGy × 2 doses) B6 CD45.1 recipients. Injected recipients were placed on Baytril-medicated water for 2 weeks after irradiation. Serum was collected at 1, 2, 4, and 6 weeks after cell transfer and chimeras sacrificed at 6 weeks to quantify splenic GCs.

### Flow cytometry

Splenocytes were harvested into ice-cold complete RPMI 1640 media (with 10% FBS, 1% pen/strep, 1% sodium pyruvate, 1% HEPES, 1% GlutaMAX and 0.1% β-ME). Cells were incubated with RBC lysis buffer (ACK) for 2 minutes, followed by quenching using the RPMI 1640 media, and then filtered through 40-μm cell strainers. Single-cell suspensions were stained with fluorescence-labeled antibodies for 20 minutes in the dark at 4°C. For the intranuclear staining True-Nuclear (BioLegend) was used. Samples were acquired using LSRII flow cytometer and LSRFortessa (BD BioSciences) and analyzed by FlowJo software (Tree Star, Inc.).

### Antibody ELISA

For autoantibody detection, 96-well Nunc-Immuno MaxiSorp plates (Thermo Fisher Scientific) were coated with 5 μg/ml Sm/RNP (ATR01-10; Arotec Diagnostic Limited) at room temperature for 2 hours. Plates were then blocked for 1 hour with 1% BSA in PBS and then followed by incubation with serially diluted serum in 1% BSA and 0.05% Tween in PBS for 2 hours. Subclass-specific IgG2a antibodies were detected using goat anti-mouse IgG2a-HRP at 1:2000 dilution (SouthernBiotech). Peroxidase reactions were developed using OptEIA TMB substrate reagent set (BD Biosciences), and reactions stopped using sulfuric acid. Absorbance was measured at 450 nm using a SpectraMax i3X microplate reader (Molecular Devices). Total serum IgG2a was measured using a sandwich ELISA kit (EMIGG2A; Thermo Fisher Scientific), as per the manufacturer’s instructions.

### B cell activation and CRISPR/Cas9 gene editing

CRISPR-mediated deletion of *Prdm1* (encoding BLIMP-1) or *Rosa26* (control locus) was performed as described (25). Briefly, splenocytes were harvested from B6 (WT) or 564Igi donors into ice-cold PBS (with 2% FBS, and 1mM EDTA). Splenocytes were filtered through 40-μm cell strainers (Corning) and enriched using EasySep Mouse B Cell Isolation Kit (19854; StemCell Tech) as per the manufacturer’s instructions. Enriched cells were cultured and activated for 2 days at a cell density of 2.5 × 10^6^ cells/mL in IMDM media (with 10% FBS, 1% pen/strep, 0.1% β-ME, 100ng/mL murine multimeric CD40L (AG-40B-0020-C010; AdipoGen) and 1μg/mL rat α-mouse CD180 (562191; BD Pharmingen)) at 37°C. After 2 days of incubation, *Rosa26* or *Prdm1* single guide RNA (sgRNA; Synthego) was complexed with Alt-R S.p. Cas9 Nuclease V3 (1081059; IDT) at a molar ratio of 3:1 in electroporation buffer (EPB-1; MaxCyte) at room temperature for 20 minutes. After 10 minute RNP incubation, B cells were harvested and resuspended in electroporation buffer at a concentration of 1 × 10^8^ cells/mL. The RNP mixture was then added, with the final Cas9 concentration of 1.25 μM, and electroporation was performed using ExPERT GTx (MaxCyte). Following electroporation, cells were resuspended in IMDM media (now with the addition of 10 ng/mL murine IL-4 ((AF)-214-14; PeproTech)) at a concentration of 4× 10^6^ cells/mL and allowed to recover at 37°C for 2 hours prior to adoptive transfer. For pilot experiments involving adoptive transfer of WT and 564Igi activated B cells, the CRISPR/Cas9 electroporation steps were omitted. 4-6 hours after reconstitution of lethally irradiated B6 CD45.1 recipients with B6 CD45.1 BM (see above), 5 × 10^6^ cells in vitro activated B cells or CRISPR/Cas9 edited B cells were transferred into recipient mice by tail vein injection.

### Statistical analysis

Statistical significance was calculated using two-tailed unpaired Student’s *t* test, or ANOVA as indicated in each figure by Prism (GraphPad) software Version 10.0. The P values were considered significant when *P* < 0.05 (*), *P* < 0.01 (**), *P* < 0.001 (***), and *P* < 0.0001 (****). No statistical tests were used to predetermine sample size.

## Data Availability Statement

Data are available in the article itself. The original data are available from the corresponding author(s) upon reasonable request.

## Acknowledgements

We thank Shari Cho, Karen Sommer, and Aesha Vakil for laboratory management.

## Author contributions

Danny Kim and Shaun Jackson conceptualized the research project and wrote the manuscript. Kristy Chiang, Andrea D. Largent, and Sivasankaran Munusamy Ponnan performed experiments and analyzed data. Ragan Pitner, Brock J. McKinney, Richard James, and David J. Rawlings assisted with design of experimental models and provided technical assistance for the CRISPR/Cas9 gene editing experiments. All authors reviewed and commented upon the manuscript.

